# Bidirectional modulation of pain by neurofeedback: Preliminary findings with fMRI at 7T

**DOI:** 10.64898/2026.05.11.724199

**Authors:** Konstantin Demin, Jun Seo Hwang, Wonyi Che, Dongho Kim, Wani Woo, Hakwan Lau, Vincent Taschereau-Dumouchel

## Abstract

Previous brain decoding studies indicate that an individual’s pain experience can be robustly predicted from distributed patterns of brain activity. Two brain decoders have notably been associated respectively with the nociceptive and cognitive aspects of pain experience, the Neurologic Pain Signature (NPS) and the Stimulus-Intensity Independent Pain Signature (SIIPS). Yet, we still do not know if these brain patterns are also causally related to pain experience. To evaluate this possibility, we used high-field (7-Tesla) fMRI to test whether humans can alter their pain experience by bidirectionally modulating their pain-related brain activity in decoded neurofeedback paradigm. In a double-blind design, participants were trained to up- and down-regulate the NPS or the SIIPS. Our results indicate that participants can achieve bidirectional control of both signatures. NPS expression reliably increased during pain stimulation and covaried with both stimulus intensity and subjective ratings. In contrast, SIIPS expression did not show consistent stimulus-locked effects in the primary analyses. Importantly, reduction in pain rating was specific for SIIPS-training, whereas NPS has failed to show any consistent behavioral effect. Based on these preliminary findings, we hereby preregister a follow-up study, with specified rationale, hypotheses, experimental design, and analysis protocols.

## 1 Introduction

Pain disorders represent a major public health concern worldwide, contributing substantially to reduced quality of life. Critically, safe and efficient treatments are often unavailable, leading many individuals to live with chronic symptoms [1–3]. Recent neurocognitive studies indicate that pain is a multifaceted experience resulting from the interaction between nociceptive input and higher-order psychological processes [4– 6] associated with widely distributed patterns of brain activity [6–8]. However, while these findings have substantially advanced our understanding of pain processing, the causal link between such brain activity patterns and subjective pain experience still remains unclear.

Decoded functional magnetic resonance imaging (fMRI) neurofeedback provides a powerful framework for testing such causal associations [9, 10]. More specifically, decoded neurofeedback allows reinforcing specific multivoxel patterns and measuring how they affect various behavioral, physiological and self-reported outcomes. This approach is thought to operate through reinforcement learning in the brain, where rewarded neural states are gradually strengthened through repeated and contingent feedback [10, 11]. Importantly, throughout the intervention, participants remain unaware of the nature and purpose of the intervention [9, 10]. As such, decoded neurofeedback can be conducted in double-blind experimental designs allowing testing the causal link between specific neural target and associated outcomes [12].

Decoded neurofeedback is particularly promising for the study of pain as well-validated multivoxel decoders already exist, which resemble brain-wide patterns with weights assigned to specific voxels allowing to estimate pain-related states in new data. For instance, it is possible to decode both nociceptive stimulation (Neurologic Pain Signature (NPS); [6]) and the higher-order psychological and contextual factors contributing to pain experience (Stimulus Intensity Independent Pain Signature (SIIPS); [13]). Each of these decoders have been extensively shown to generalize across individuals and experimental contexts [14–17], making them interesting targets for neurofeedback.

In a recent proof-of-concept study, we demonstrated that participants can indeed selectively downregulate whole-brain SIIPS expression in a double-blind design [18]. More specifically we showed that SIIPS expression could be down-regulated independently from NPS expression, suggesting that the two decoders might indeed be selectively modulated using decoded neurofeedback. Furthermore, the downregulation of SIIPS was associated with the reduction in subjective pain ratings following the intervention. However, these proof-of-principle findings did not fully address how modulating SIIPS and NPS could impact pain experience as only the SIIPS decoder was targeted by the intervention.

Here, we propose to expand on our previous findings and investigate, in a proof-of-principle study, if the SIIPS and NPS decoders are causally linked to pain perception. Notably, we propose to expand on our previous approach by using high-field fMRI. So far, few neurofeedback studies have been conducted at 7-Tesla [19] but this imaging method may present a great improvement as it presents a better signal-to-noise ratio [20].

To anticipate, we found that participants can reliably up- and down-regulate both signatures. Furthermore, we found that SIIPS-training resulted in reduction in pain rating whereas NPS-training didn’t produce significant behavioral effects. Based on this pilot data we further preregister a follow-up study design and hypotheses to be tested in a broader experiment.

## 2 Methods

### 2.1 Participants

8 healthy adult participants were recruited from the local university community (5 females, 3 males, mean age = 23.6 *±* 2.36 95% CI years) for the experiments. Inclusion criteria were: age between 18 and 45 years, normal or corrected-to-normal vision, no current or past diagnosis of neurological, psychiatric, or chronic pain disorders, no use of psychotropic or analgesic medications, and no contraindications to high-field MRI. All participants provided written informed consent before participation. The study protocol was approved by the relevant institutional review board of Sungkyunkwan University (SKKU IRB 2025-02-085) and conducted in accordance with the Declaration of Helsinki.

As this study is explicitly treated as a pilot, the sample size was determined based on feasibility constraints and prior decoded neurofeedback studies in pain, rather than formal power calculations. The pilot data are intended to inform effect size estimates and stopping rules for a subsequent larger-scale confirmatory study.

### 2.2 Magnetic resonance imaging parameters and apparatus

Functional images were acquired using a T2*-weighted gradient-echo echo-planar imaging (EPI) sequence on a 7-Tesla Siemens MAGNETOM Terra scanner with a 32-channel head coil at the Center for Neuroscience Imaging Research, Institute for Basic Science, Suwon, South Korea. Acquisition parameters were as follows: repetition time (TR) = 1400 ms, echo time (TE) = 21.0 ms, flip angle = 61°, voxel size = 1.5 × 1.5 × 1.5 mm, field of view (FOV) = 210 × 210 mm, 92 axial slices, phase-encoding direction = posterior to anterior, multiband acceleration factor = 4, GRAPPA acceleration factor = 2, partial Fourier = 6/8, bandwidth = 1984 Hz/pixel. Slice orientation was set obliquely to the AC-PC line (approximately -15° to axial).

At the beginning of the first session, structural T1-weighted images were acquired. A 3D Magnetization-Prepared Two Rapid Acquisition Gradient Echoes (MP2RAGE) sequence was used with the following parameters: TR =3700 ms, TE = 1.5 ms, inversion times (TI1/TI2) = 1000/3200 ms, flip angles = 4°/4°, voxel size = 1 × 1 × 1 mm, FOV = 210 × 210 mm, 192 sagittal slices, phase encoding direction = anterior to posterior, bandwidth = 370 Hz/pixel, GRAPPA acceleration factor = 3. Background denoising was disabled, and the standard MP2RAGE UNI image was used as the T1-weighted anatomical reference for subsequent preprocessing.

To correct for susceptibility-induced geometric distortions in the functional data, a pair of gradient-echo EPI volumes with reversed phase encoding directions, one with anterior-to-posterior and one with posterior-to-anterior, was acquired before each functional run. These images were acquired with the same pulse sequence as the functional scans. For each phase encoding direction, four volumes were acquired in an interleaved slice order, and the final volume of each pair was used for distortion correction.

### 2.3 Stimuli Presentation and Hardware

Visual stimuli were displayed on a screen behind the scanner and observed through a mirror attached to the head coil. The refresh rate and spatial resolution of the monitor were 60 Hz and 1600 × 1000 pixels, respectively. The effective viewing distance was 104 cm. Visual stimuli were presented using a separate working station (display PC) synchronised with the real-time PC.

The real-time PC was a desktop stationary computer running on Ubuntu 22.04 with Intel Core i9-14900K CPU, NVIDIA GeForce RTX 4090 (24 GB VRAM) GPU, 64 GB DDR5-5600 RAM, and Samsung 990 PRO NVMe SSD (2 TB). The display PC was a Windows 11 Pro laptop with 12th Gen Intel Core i5-12450H, NVIDIA GeForce RTX 4060 laptop (8 GB VRAM) GPU, 16 GB DDR5-4800 RAM, and Samsung SSD (1 TB).

### 2.4 Real-Time Preprocessing and Decoding

Real-time preprocessing was implemented using a GPU-accelerated SPM-based (SPM12 was used [21]) pipeline adapted from the ATR toolbox (https://bicr.atr.jp/decnefpro/software/) and the prior pain decoded neurofeedback study [18]. The experimental code was run in MATLAB R2024b (MathWorks, Natick, MA, USA) with the Psychophysics Toolbox Version 3.0.19 [22] using the real-time PC station described above. As functional images were acquired, they were transferred to a dedicated real-time processing workstation and subjected to one of the following protocols:

#### 2.4.1 Transformations generation (EPI->T1->MNI)

This step is conducted before the real-time procedures in an “offline” regimen, taking approximately 90 seconds at the start of each run, aiming to generate a transformation file that will be further applied in the real-time protocol (Table 1). During the Resting State run (which is conducted first on each day), some additional steps are included, taking about 10 minutes in total.

**Table 1.**
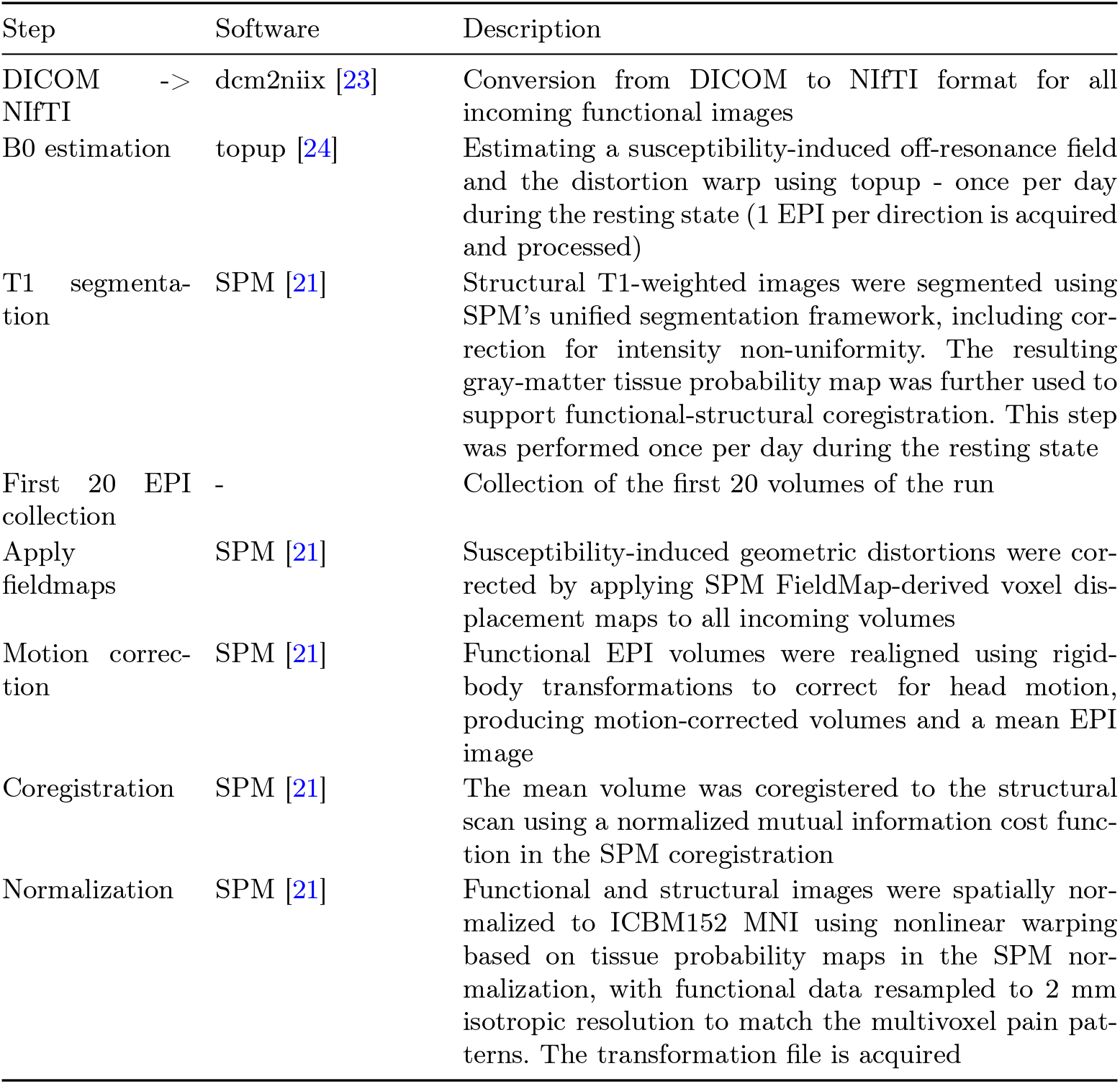
Transformations generation pipeline.

#### 2.4.2 Real-time procedure

This is the main analyses step, in which the precomputed transformations are applied to each incoming functional volume and neurofeedback scores are calculated from the decoder output (Table 2). These operations were performed online throughout the run so that participants could receive feedback on their current brain state with minimal delay.

**Table 2.**
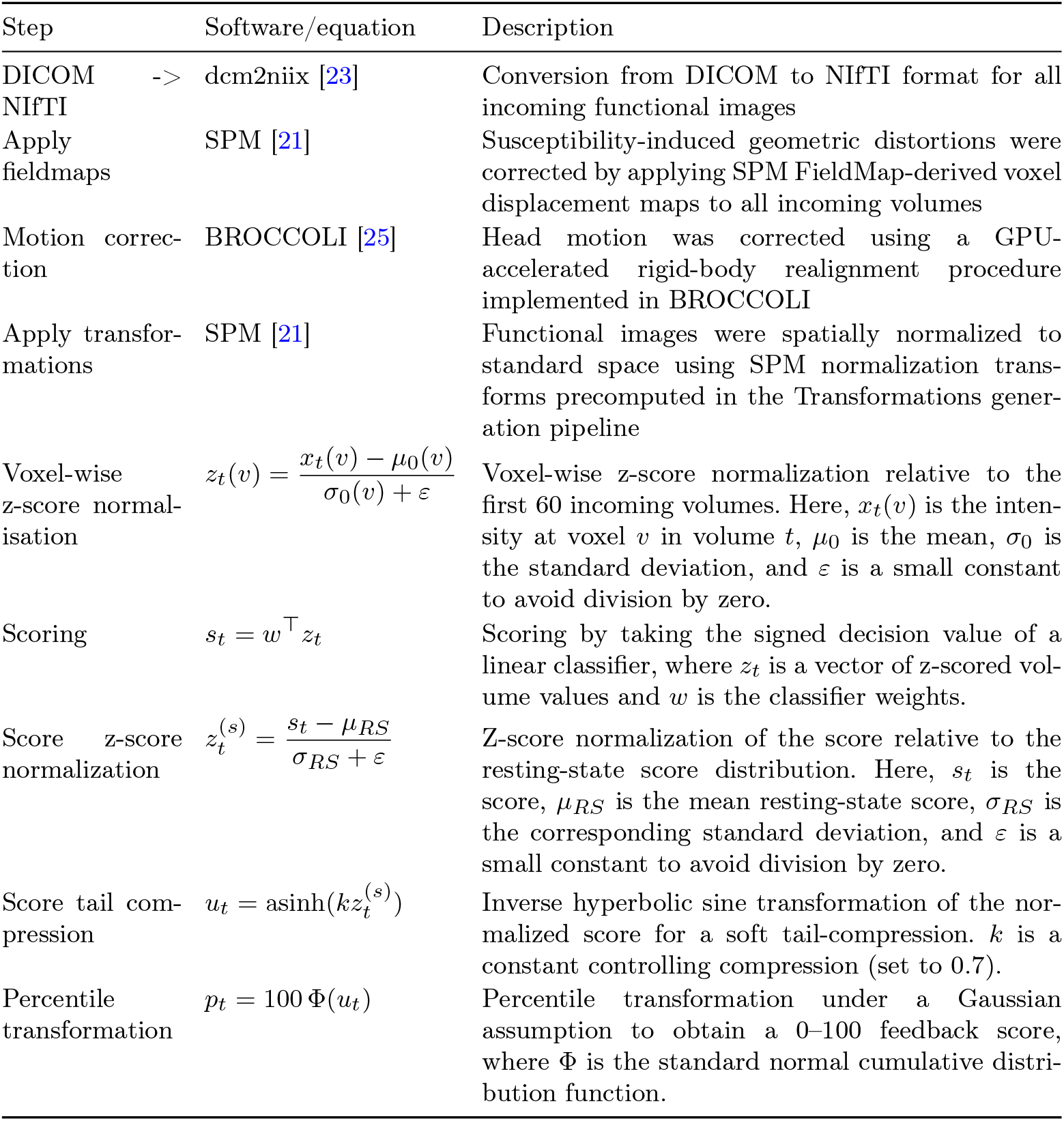
Real-time procedure pipeline.

Together, this pipeline converted each newly acquired volume into a standardized decoder score that could be compared across runs and days. The final percentile-transformed value was then used as the feedback signal shown to participants during regulation, providing a compact summary of ongoing pain-related brain activity before moving to the broader experimental design.

### 2.5 Experimental Design Overview

Unlike previous work [18], here we implement a within-subject design in which each participant is trained to modulate the same pain-related pattern in two opposing directions (up- or down-regulation) within the same day, allowing direct assessment of bidirectional causal effects on subjective pain within individuals (see Fig. 1).

**Fig. 1.**
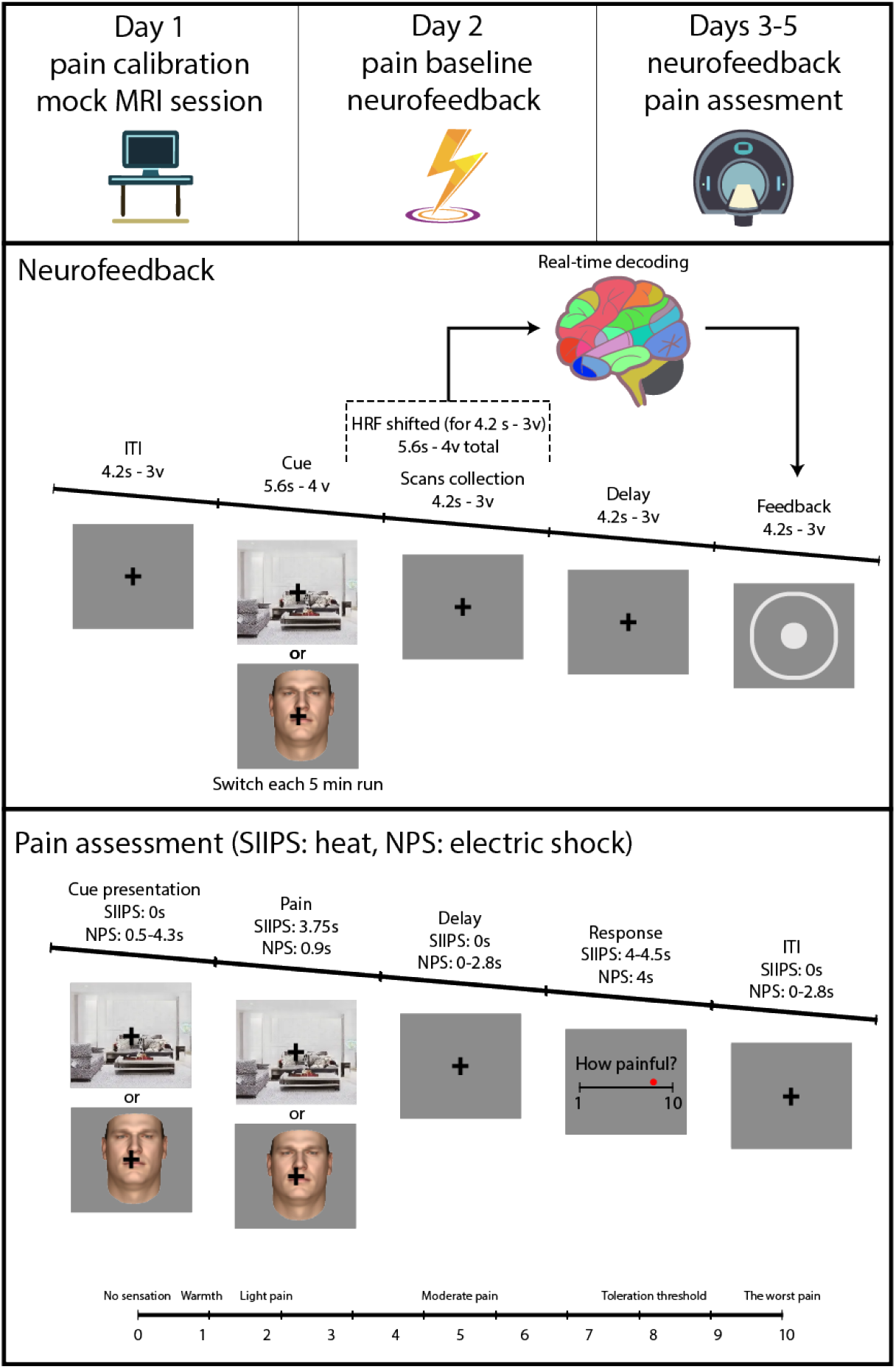
Task Designs. Top panel: The Decoded Neurofeedback protocol used in the experiment. Bottom panel: Pain Assessment in the SIIPS group (heat stimulation) and NPS group (electrical stimulation). s - the duration of the event measured in seconds; v - the duration of the event measured in volumes; HRF - Hemodynamic Response Function. On each neurofeedback trial, participants viewed one of the visual natural cues (Fig. 1 - Cue; Supplementary Fig. 1) - a face (synthesised in Basel Face Model 2019, [26]) or a scene (from a continuous space of Scene Wheels, [27]) - while their ongoing brain activity was recorded and analysed in real-time. In SIIPS group (decoder up- or down-regulation) cue was changed pseudorandomly each run (conditions balanced out), whereas in NPS group, all the runs of one condition were grouped together in a blocked sequence (Table 3).

The study’s global design included

**Table 3.**
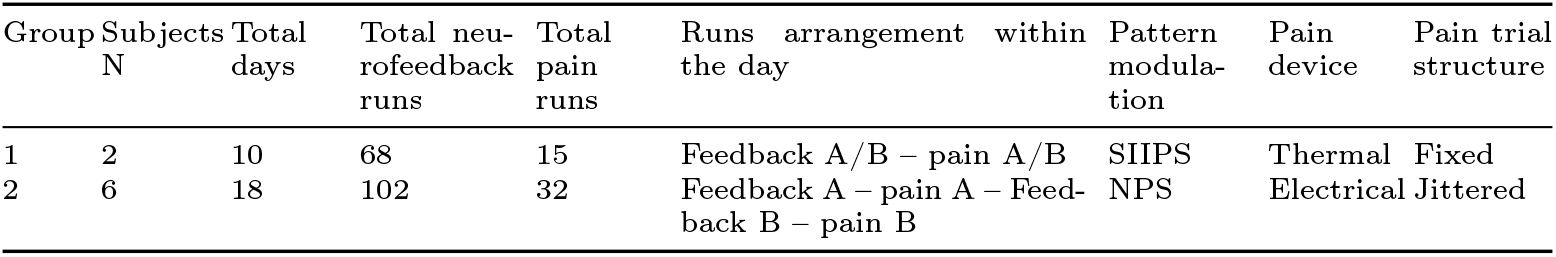
Key differences between the two groups.

- Day 1. A pain calibration session outside the scanner was followed by a pain session (including 32-40 trials), a 10-minute mock MRI session, and a second pain session.
- Day 2. The pain baseline session inside the scanner consisted of a 10-minute resting state run, 2 pain sessions (total of 64-80 trials), followed by 1-3 runs of neurofeedback training (14 trials each) per condition.
- Training days (3-5, depending on the subject). Main neurofeedback sessions inside the scanner, consisting of 10 minutes resting state run, followed by 1-3 runs of neurofeedback training (14 trials each) per condition, followed by 1-2 pain runs (32-40 trials each).

The target decoder was either the Stimulus Intensity Independent Pain Signature (SIIPS, n=2) or the Neurologic Pain Signature (NPS, n=6), depending on the experimental group (see Table 3). Participants were blind both to the identity of the decoder and the direction of modulation. Experimenters were blind to the direction of modulation. We assessed the effect of neurofeedback on painful stimulations (delivered during Pain runs).

Given the pilot nature of the study, one of its key aims was to inform further preregistration decisions. For that reason, slightly different experimental designs were used in different sessions. Most of them were devoted to finding a pain delivery and assessment protocol that would be sensitive experimentally and suitable logistically. Table 3 outlines key experimental protocols.

The SIIPS and NPS groups were also different in terms of training days (SIIPS: subjects n=2; NPS: subjects n=6), pain device used (SIIPS - thermal, NPS - electrical), and pain trial structure (see pain assessment procedure below for the exact timings). The groups were also different in the neurofeedback and pain runs arrangement within the training days. More specifically, the SIIPS group was first administered all neurofeedback runs for all conditions, and then the two pain runs. The NPS group was first administered one neurofeedback condition (up- or downregulation), followed by a pain run, and then took part in the second neurofeedback condition and its associated pain runs. For the smaller differences varying between the sessions (i.e., in event timings) we report them as ranges, where appropriate.

### 2.6 Pain decoders

We used two different brain decoders for the neurofeedback training and behavioral pain testing: The Neurologic Pain Signature (NPS [6]) is a multivariate fMRI brain pattern developed to quantify neural responses that reliably track subjective pain intensity driven by direct nociceptive input. It consists of distributed voxel weights across regions strongly involved in processing noxious stimuli (e.g., insula, anterior cingulate, thalamus). The decoder was trained on different datasets of evoked painful stimulation (heat, mechanical, electrical) and is minimally responsive to other non-painful aversive stimuli.

Stimulus-Intensity Independent Pain Signature (SIIPS [13]) was developed to capture cerebral contributions to pain that are not explained by nociceptive stimulus intensity or the NPS response (by regressing their effects out). This decoder is a whole-brain pattern that includes substantial weights in prefrontal, limbic, and other regions implicated in cognitive, evaluative, and motivational aspects of pain.

### 2.7 Decoded Neurofeedback Procedure

Each Decoded Neurofeedback run consisted of 14 trials including: (1) the cue presentation (i.e., a face or a living room), (2) a period of shifted scan collection to account for the hemodynamic response function delay, (3) a waiting period for volumes transfer and analysis, (4) feedback (communicated through the diameter of the feedback circle, see Fig. 1A) and (5) an inter-trial interval (Fig. 1). The duration of each event and trial was completely aligned with the scanner volume acquisition time (TR), avoiding the necessity for data interpolation or exclusion. All events and preprocessing steps were applied identically across upregulation and downregulation conditions.

No explicit cognitive or imagery strategies were suggested. Participants were instructed only to “maximize the score” and “increase the circle size”, consistent with reinforcement learning-based neurofeedback paradigms [9, 10]. An additional monetary reward was paid to subjects whose daily performance was better than that of the previous subject/session.

To account for HRF during scoring, only the volumes that span a period starting 4.2 seconds or 3 volumes following the cue presentation were analyzed (Fig. 1 - Scans Collection, HRF shifted). Scans were then collected for another 4.2 seconds, and the 4.2-second delay was introduced afterwards to allow for the data transfer from the scanner PC to the real-time PC and the analysis (Fig. 1 - Delay).

At the end of each trial, feedback (Fig. 1 - Feedback) was presented visually as a circle, the size of which was proportional to the decoded expression of the target multivoxel pattern z-normalized to the Resting State data. For upregulation trials, higher decoder expression resulted in larger feedback, whereas for downregulation trials, lower decoder expression was rewarded. To keep subjects engaged, additional cues were provided when decoder expression exceeded a 70th percentile threshold relative to a resting-state baseline distribution.

Additionally, to keep subjects looking and focusing on the presented cues, a small attention and working memory task was implemented, where they had to count how many times a cue picture was different from the aim (Supplementary Fig. 1). Cue had a 2.5% chance to be different from the aim on each trial, but the cumulative number of presented distractors couldn’t be greater than 3. At the end of each run, subjects had to report how many times the distractors were presented. Responses were collected using a trackball device.

### 2.8 Real-time processing of feedback

A 430 TR long (about 10 minutes) resting state run was performed at the beginning of each session to determine the distribution of corresponding pain-decoder (NPS or SIIPS) expression, similar to [18]. The volumes were analyzed and processed similarly to the real-time neurofeedback data, and the resulting distribution was further used to z-score the neurofeedback data relative to this resting state pain-decoder expression distribution.

Because the linear decoder produces values in arbitrary units determined by voxel scaling and model weights, their absolute magnitude is not directly interpretable. By estimating the mean and variance of decoder expression during resting state and z-normalizing ongoing decoder values relative to this distribution, we converted raw classifier outputs into standardized deviations from each participant’s day-specific baseline.

This normalization step also served three other purposes. First, it defined decoder expression relative to within-day variability, enabling meaningful comparison across sessions and individuals. Second, it minimized the influence of scanner drift, global signal changes, and day-to-day variance in overall signal scaling. Third, it enabled feedback to be expressed as percentile deviation from spontaneous decoder expression, thereby stabilizing the reinforcement contingency across training days. All reported decoder values reflect this resting-state-normalized scale.

### 2.9 Pain calibration

The pain calibration task was conducted before any other pain tests and consisted of 12 stimulations and was designed akin to heat pain calibration studies [28]. Briefly, 3 stimulations with a predetermined intensity values (43.4, 45.4, 47.4°C or 15, 20, 25 mA) are first delivered in random order. Based on these responses, a linear regression model is fitted to predict the pain ratings from the current. This linear regression was updated continuously based on further participant responses. Each time, three intensity values are estimated corresponding to 30, 50, or 70 visual analog scale (VAS) ratings, where 0 indicated “no pain” and 100 indicated “worst pain imaginable” (Fig. 1; [6, 13]). The next stimulation is selected pseudo-randomly (ensuring no repetition) between those three estimations. Based on the final trial linear regression data, we estimated four intensity values equally spaced between 30 and 70 VAS estimation (30+13.(3)*n) that were used for the rest of the study. The maximum °C estimation for thermode couldn’t exceed 48°C for any VAS rating, and the mA estimation couldn’t exceed 40 mA.If the estimated intensity exceeded the limit, linear regression outputs were automatically capped to this value.

### 2.10 Pain assessment procedure

During pain assessment runs, thermal stimulation was applied at four individually calibrated intensities (equally spaced between 30 and 70 ratings). Participants rated perceived pain intensity following each stimulation using the Visual Analog Scale (VAS). All responses were collected using a trackball device. Pain ratings collected during fMRI pre- (day 2) and post- (days following day 2) neurofeedback sessions constituted the primary behavioral outcome measure.

The exact pain assessment sessions structure varied depending on the group (Table 3, Fig.1). Briefly, in the SIIPS group, thermal stimulation was administered during the neurofeedback cue presentation. Stimulation lasted for a total of 3.75 s, including heating (0.25s) and cooling (0.5s) periods. After each trial, VAS rating response were collected in approximately 4-4.5s. No inter-trial intervals or other waiting periods were included in the trials. Each trial lasted up to 8.25s, and 32 trials were acquired, separated into two sessions. Cue presentation conditions (decoder up- or down-regulation) were randomly switched in blocks of 4 trials. On the pretraining day, both pain assessment sessions were applied sequentially before the first neurofeedback session, whereas for the rest of the days, both pain assessment sessions were done at the end of the day, following all the neurofeedback training runs.

In the NPS group, a cue was presented for 0.5 - 4.3s, followed by a short 0.9s 3 Hz train of electrical stimulation (Fig. 1). The rating response event (4s) was preceded by a 0-2.8s jittered delay (without a cue presentation). Finally, trials were separated by the inter-trial intervals. Each trial lasted up to 14.8s, and 40 trials were acquired, separated into two sessions. Similar to SIIPS group on the pretraining day, both pain assessment sessions were applied sequentially before the first neurofeedback session. However, for the rest days, pain assessment sessions were grouped with the corresponding neurofeedback runs by condition. All the runs of one condition of the neurofeedback were performed first, followed by the pain assessment run, consisting only of the same condition trials. Then, the other condition’s neurofeedback and pain assessment run follows.

Similarly to the neurofeedback runs, to keep subjects looking and focusing on the presented cues, a small attention and working memory task was implemented, where they had to count how many times a cue picture was different from the aim (Supplementary Fig. 1). Cue had a 2.5% chance to be different from the aim on each trial, but the cumulative number of presented distractors couldn’t be greater than 3. At the end of each run, subjects had to report how many times the distractors were presented.

### 2.11 Pain stimulation

Thermal stimulation was delivered to the distal lateral lower leg, proximal to the lateral malleolus (akin to [29]), using an fMRI-compatible 16 × 16 mm ATS thermode (Medoc, Israel) passed into the scanner room through a waveguide. Stimuli temperature ranged from 39.2 to 48°C with baseline temperature of 32°C and an incremental step of 0.2°C. The thermal stimulation had a duration of 3.75 s: 0.25 s ramp-up (heating), a 3-s plateau, and a 0.5-s ramp-down (cooling). The ramp speeds were adjusted to fit specific windows independently of the temperature, with heating rates being 28.8–72°C/s and cooling rates 14.4–36°C/s.

Electric pain stimulations were delivered using the DS8R Biphasic Constant Current Stimulator (Digitimer Ltd., UK) with an fMRI-compatible D185-HB4 Digitimer Ltd., UK) cable passed into the scanner room through a waveguide and via pre-wired disposable electrocutaneous stimulation electrodes (round, ≈ 20 mm diameter, adhesive hydrogel contact, cloth-backed). Electrodes were attached to the left arm over the distal (inferolateral) deltoid region near the deltoid tuberosity (lateral humerus), with the distance between electrode cores being 2.5 cm and the distance between electrode gelpads being 0.5 cm. The electrode position was chosen based on the internal testing and the fact that upper arm stimulation may result in fewer involuntary muscle contractions and better discriminability [30]. Biphasic, charge-balanced stimulation with alternating leading polarity was applied as a train of three pulses, with 200-µs phase duration and a 50-µs interphase interval, separated by ≈ 300 ms intervals. Applied current ranged from 1 to 40 mA.

### 2.12 Data analysis

#### 2.12.1 Decoder induction

Decoder induction was estimated during the neurofeedback sessions. To quantify the brain decoders’ relative activity, the decoder output (i.e., predicted scores from the SIIPS or NPS) were z-normalized relative to this day’s resting-state decoder distribution and used for further analysis (also see score estimation section).

During each trial, the decoding window started 3 volumes after cue presentation (to account for HRF) and lasted for 4 volumes (Fig. 1). Data within the decoding window were pre-processed (see Real-time processing of feedback) and the decoder output was analyzed both at the run and day levels. First, we computed a single run-level mean (14 trials, scoring volume blocks) for each combination of subject × day × run × decoder (SIIPS or NPS) × condition (Up- or Down-regulation). A linear mixed-effects model (MATLAB fitlme) was used to estimate decoder, condition, and decoder×condition fixed-effects, as well as subject:day nested random effects on the brain decoder expression mean using omnibus tests (F-tests):

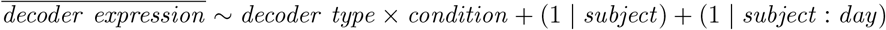

Second, within each day, we computed the induction difference (delta) between the up and down conditions for each decoder, defining the within-day induction effect as:

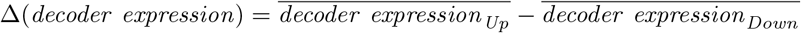

Then, we tested whether the delta differed from zero within each decoder using intercept-only mixed models or whether deltas differed between decoders using the following linear mixed-effects models (reporting fixed-effects using omnibus tests (F-tests)):

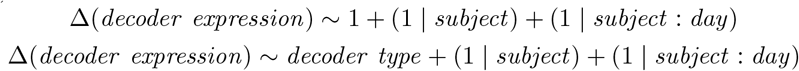

Furthermore, to investigate the dynamic changes in decoder expression within a trial, a fixed cue post-onset window spanning 15 volumes was used. Then, for each decoder (SIIPS or NPS) separately, we modeled the event-locked time course (linear mixed-effects model) using volume sequential number as a categorical factor, condition as a categorical factor (Up vs. Down), and their interaction (reporting fixed-effects using omnibus tests (F-tests)):

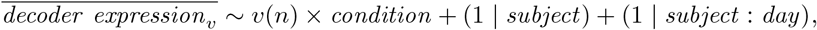

where *v* denotes the volume and *v*(*n*) denotes the volume number.

When a significant volume × condition interaction was observed in the mixed-effects model, post-hoc contrasts were performed to test the simple effect of condition (Up vs. Down) at each post-cue volume. These contrasts were computed as linear combinations of fixed-effect coefficients from the same fitted interaction model.

For the tests estimating the dynamic effects performed in this section, p-values were adjusted using Benjamini-Hochberg false discovery rate (BH-FDR) corrected across all omnibus time course tests for both decoders. We report FDR-adjusted q-values in the plot annotations.

#### 2.12.2 Correspondence between pain decoder predictions and pain ratings

To validate that the pain decoders track pain ratings, we collected the data of both groups (Table 3) during pain assessment runs and pain VAS ratings. The decoder outputs (both SIIPS and NPS) were z-normalized relative to this day’s resting-state decoder distribution (also see score estimation section) and treated as dependent variables in all subsequent analyses.

Decoder outputs were analyzed using linear mixed-effects models (LMEs) to characterize pain-induced temporal dynamics and decoder effects. We fit a pain-event stimulation-locked HRF-convolved regressor using an impulse train of ones at each onset volume *t*. Then we fit a separate linear regression model:

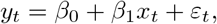

where *x*_*t*_ is the HRF-event-locked regressor, *y*_*t*_ is the decoder prediction, and *β*_1_ is the estimated coefficient associated with the convolved regressor *x*_*t*_.

Next, to obtain subject-level estimates we used a hierarchical aggregation procedure. The run-level *β* coefficients obtained before were averaged within each experimental session (subject and day) to account for the non-independence of runs acquired on the same day. Finally, day-level estimates were averaged within each subject, yielding a single summary value per subject. Group-level effects were assessed by testing subject-level mean coefficients against zero using one-sample t-tests.

To further characterize event-locked coupling between reported pain intensity and decoder expression, we computed lagged within-run correlations between trial-wise pain ratings and decoder values sampled at fixed temporal offsets relative to pain onset. For each run and lag, we computed a Pearson correlation *r* between ratings and lagged decoder values. Correlations were then Fisher-z transformed, *z* = atanh(*r*), prior to aggregation.

To obtain one estimate per subject, we averaged Fisher-z values hierarchically (run:day:subject), and group-level inference at each lag was further used to test whether the subject-level Fisher-z mean differed from zero using two-sided one-sample t-tests. Multiple comparisons across lags were controlled within each decoder using Benjamini-Hochberg FDR (BH-FDR), and results are reported as q-values.

#### 2.12.3 Decoder induction success during pain assessment

This analysis follows from the preceding one. Here, we aimed to understand whether experimental conditions affected the association between decoder output and pain ratings. As such, we performed the same analysis as in the previous section, but this time we analyzed the groups (Table 3; SIIPS and NPS) separately. The decoder outputs were z-normalized relative to a given day’s resting-state decoder distribution (also see score estimation section) and treated as dependent variables in all subsequent analyses.

Decoder outputs were analyzed using linear mixed-effects models (LMEs) to characterize pain-induced temporal dynamics and condition effects. For each run × condition (Up- or Down-regulation) pair, we fit an HRF-convolved model as described above.

At the run-level, each decoder (SIIPS, NPS) and event-regressor run-level *β* estimates from upregulation runs were compared to those from downregulation runs using a two-sample Welch t-test. Additionally, for each condition separately (Up- or Down-regulation), estimates were tested against zero using a two-sided one-sample t-test. Mean estimates and 95% confidence intervals were computed across runs within each condition.

To assess within-day effects, run-level beta estimates were first averaged within each subject × day × condition. For each subject × day unit, a within-day contrast was then computed as:

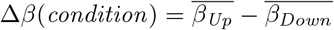

These daywise Δ*β*(*condition*) values were computed separately for each decoder and event-locked-regressor. Daywise deltas were tested against zero using a one-sample t-test. Mean deltas and 95% confidence intervals were computed across subject × day units.

#### 2.12.4 Effects of training on pain ratings

Pain ratings were analysed at the day level to keep the inferential units comparable across subjects (including cases with only a few participants, as was the case for this pilot data). The pipeline was applied separately for SIIPS and NPS groups.

Pain data were averaged by prevs. post-training (day 2 vs. any other fMRI day) and condition (Up-vs. Down-regulation). From this, we further computed the within-day difference between the two conditions (the condition effect):

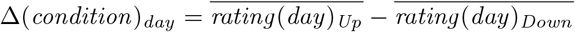

And then for each subject, we also contrasted the Δ(*condition*)_*day*_ of post-training to a corresponding Δ(*condition*) pre-training baseline as the mean Δ(*condition*)_*day*_ across all available pre days, getting baseline corrected condition effect for each day:

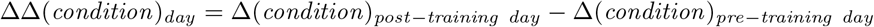

All inferential tests were performed on these day-level aggregates, with subject as a random factor. We ran the same analysis pipeline separately for the SIIPS and NPS neurofeedback datasets.

For each of these metrics, we fit one of the following linear mixed-effects (LME) models (MATLAB fitlme). We fit a linear mixed-effects model with fixed effects of training, condition, and their interaction, and a random intercept for a subject. We tested the main effects and interaction using t-tests on the corresponding fixed-effect coefficients:

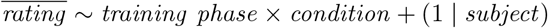

We then modeled day-level Δ(*condition*) values with training as a fixed effect and subject as a random intercept, and tested whether Δ(*condition*) differed between pre- and post-training using the t-statistic for the training effect:

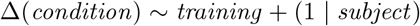

Finally, we tested whether baseline-subtracted ΔΔ(*condition*) values differed from zero using a two-sided one-sample Student’s t-test at the day level:

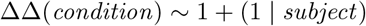

Additionally, to examine whether training effects depended on stimulus intensity, we analysed pain ratings as a function of within-day intensity rank (separately for SIIPS and NPS). First, we presented stimuli intensity as a ranked list of four pain stimulus intensities from 1 (lowest) to 4 (highest) for each subject. Then, for each subject × day × training phase × condition × pain stimulus intensity combination, we calculated their mean values. We further fit a linear mixed-effects model:

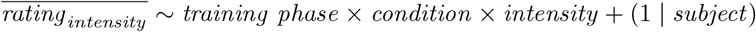

And report F-tests from the marginal ANOVA table for the main effects and interactions.

Then, analogous to the total mean rating analysis, we calculated the within-day difference between the two conditions (the condition effect Δ(*condition*)) for each intensity and Δ(*condition*)_*day*_ of post-training to a corresponding Δ(*condition*) pretraining baseline as the mean Δ(*condition*)_*day*_ across all available pre days, getting baseline corrected condition effect (ΔΔ(*condition*)) for each day × intensity pair. We then fit two linear mixed-effects models, assessing if effects for both parameters are different from zero:

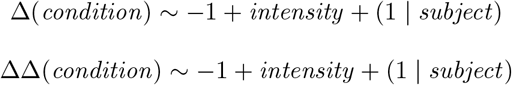

The values were controlled for multiple testing across ranks within each phase using Benjamini-Hochberg FDR (*α* = 0.05).

## 3 Results and discussion

### 3.1 Neurofeedback induction success

To assess induction effects, normalized decoder predictions during induction were analyzed using a mixed-effects model. This analysis revealed significant main effects of decoders (F(1, 166) = 7.08, p = 0.0086) and conditions (F(1, 166) = 7.51, p = 0.0068; Fig. 2 left-top panel). These results indicate that SIIPS and NPS differed in their overall induction and that upregulation runs exhibited higher decoder values than downregulation runs, indicating the success of induction. Importantly, no decoder × condition interaction was observed (F(1, 166) = 0.006, p = 0.94).

**Fig. 2.**
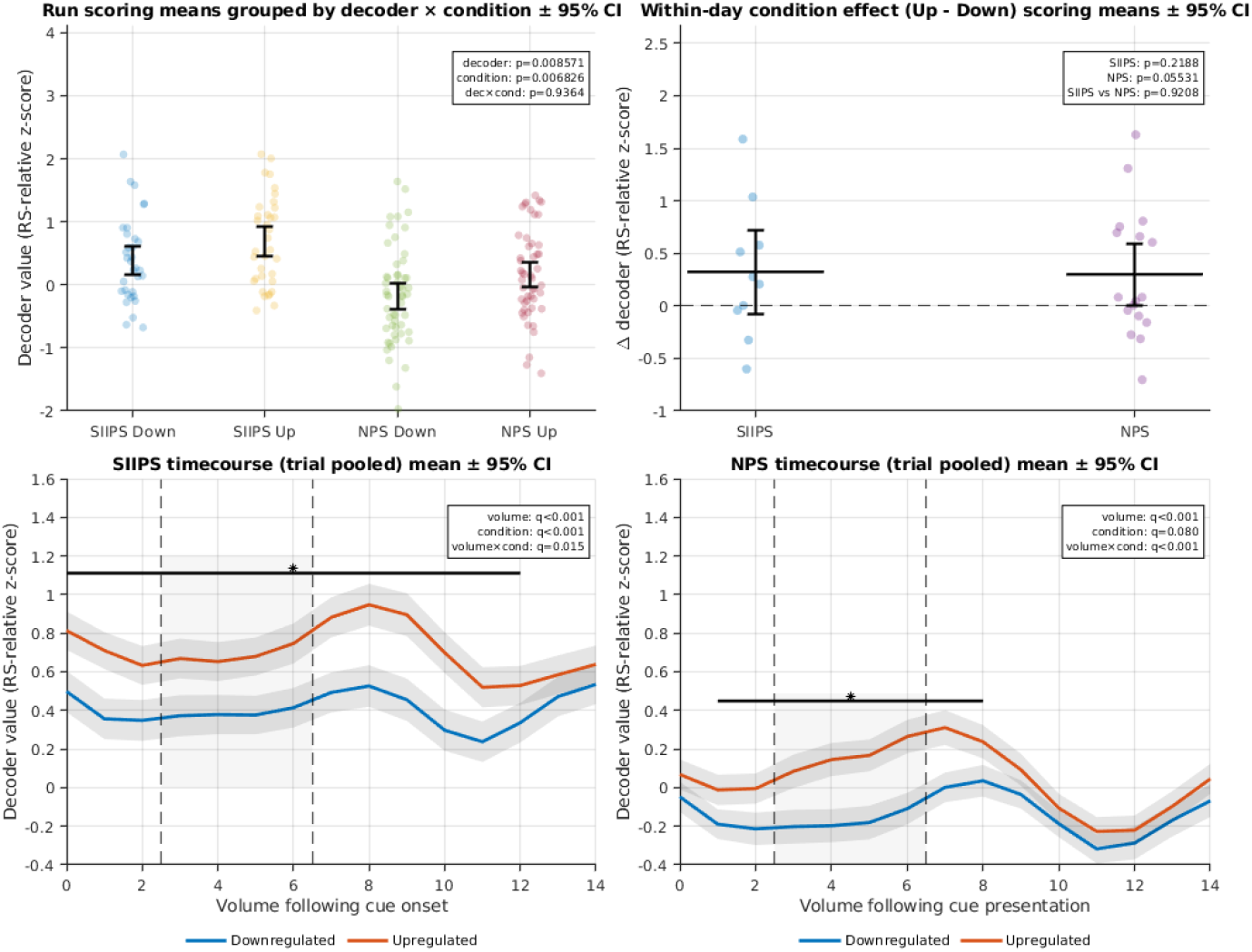
Neurofeedback-related modulation of activity (SIIPS or NPS). The top-left panel shows responses from the different experimental conditions (Down = downregulation, Up = upregulation), in terms of the averaged decoder output after each neurofeedback trial. The top-right panel is based on the same data, but shows the differences of effects between up- and down-regulation within the same day, on decoder output. Bottom panels show the dynamics of decoder output within a trial, for SIIPS sessions and NPS sessions, respectively. All decoder values are expressed as z-scores relative to the statistical distribution derived from resting-state data. The dotted regions on time course plots represent the range of volumes used for scoring. All plotted uncertainty bands and error bars reflect 95% confidence intervals. * - q<0.05 for post-hoc linear mixed model simple effect contrasts of condition × volume interaction effect (Up-vs. Down-regulation) corrected using Benjamini-Hochberg FDR.

We next examined induction effects within-day by computing, for each subject × folder × decoder, the difference between upregulation and downregulation mean normalized decoder values (Δ = Up - Down) within the same volume range. Intercept-only mixed-effects models testing whether Δ differed from zero revealed no significant within-day induction effect for either decoder (SIIPS: F(1, 9) = 1.75, p = 0.219; NPS: F(1, 16) = 4.27, p = 0.055; Fig. 2 top-right panel). However, the NPS group was very close to approaching statistical significance, even considering the relatively low power of the analysis compared to other decoded neurofeedback studies. Finally, a mixedeffects model comparing deltas between the decoders showed no difference between SIIPS and NPS (F(1, 25) = 0.01, p = 0.921).

We then analysed cue-presentation locked decoder responses separately for SIIPS and NPS subjects. SIIPS responses following neurofeedback cue presentation showed significant modulation across volumes and a significant condition effect (Up-vs. Down-regulation; Fig. 2 bottom-left panel). A mixed-effects model including postcue volume, condition, and their interaction revealed a main effect of volume (F(14, 13230) = 2.97, q = 2.22 × 10^−4^ BH-FDR), a main effect of condition (Up vs. Down; F(1, 13230) = 18.55, q = 3.34 × 10^−5^ BH-FDR), and a significant volume × condition interaction (F(14, 13230) = 2.04, q = 0.015 BH-FDR). Consistent with the significant interaction effect, post-hoc simple effect contrasts comparing Up-vs. Down-regulation at each post-cue volume identified significant condition differences at 13 of 15 volumes.

We performed the same time-course analysis for NPS decoder modulation. The full mixed-effects model also showed a strong main effect of volume (F(14, 19860) = 7.52, q = 3.07 × 10^−15^ BH-FDR) and a highly significant volume × condition interaction (F(14, 19860) = 4.18, q = 6.75 × 10^−7^ BH-FDR). Post-hoc volume-wise contrasts showed significant Up-Down differences at 8 of 15 volumes. In contrast to the SIIPS timecourse, the condition main effect did not reach significance in the interaction model (F(1, 19860) = 3.07, q = 0.080 BH-FDR), indicating that Up-Down differences were not uniform but interacted with cue presentation.

Summing up this section, subjects successfully Up- and Down-regulated their pain-related brain activity . To our knowledge, these results are the first to indicate that humans can achieve bidirectional control of these brain decoders in real-time fMRI.

### 3.2 Decoder expression during pain assessment

Across all days, NPS showed reliable positive pain coupling, whereas SIIPS failed to show any association with pain (Fig. 3). For event-locked models (event × HRF) - NPS mean *β* estimates were significantly greater than zero (t(7) = 5.895, p = 0.0006026), whereas SIIPS did not (t(7) = -0.138, p = 0.8944).

**Fig. 3.**
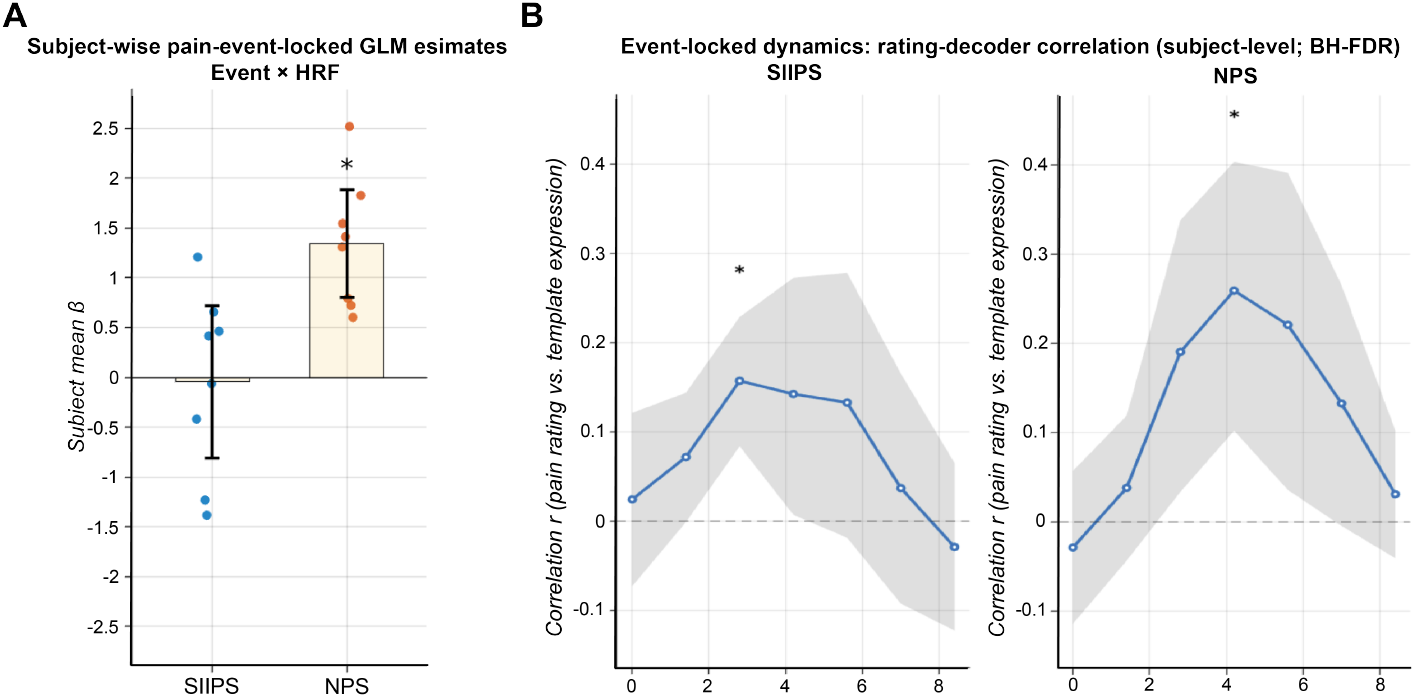
Pain encoding by the pain decoder activity (SIIPS or NPS). The panel A shows the degree to which subjects’ self-rating of pain predicted the expression of the brain pain decoders, respectively, across trials. Shown are based on data from all experimental sessions included in both groups. This degree was quantified using *β* estimates obtained from HRF-convolved pain stimulation onsets GLM regressor applied to decoder time series z-scored relative to the distribution of resting-state data (see Methods). For each subject (n = 8), run-level *β* estimates were averaged hierarchically (run - day - subject) to yield a single value per subject. Mean run-level *β* estimates represented as bars, with dots indicating individual runs and error bars denoting 95% confidence intervals. The panel B shows event-locked dynamics of rating-brain decoder coupling, i.e. the degree to which pain ratings predicts corresponding pain decoder values, at each time point after pain stimulus presentation. Mean correlation between trial pain ratings and decoder expression (z-scored relative to the statistical distribution derived from resting-state data) sampled at fixed lags after the corresponding trial pain onset, shown separately for SIIPS and NPS. Correlations were computed within each run at each lag, Fisher-z transformed, and aggregated hierarchically across runs and days to yield one value per subject. Lines denote the group mean correlation (back-transformed from Fisher-z); shaded regions show 95% confidence intervals across subjects (n = 8). Asterisks mark lags with q < 0.05 by Benjamini-Hochberg FDR following two-sided one-sample t-tests.

Additionally, we performed within-subject comparisons of the decoder that further indicated that NPS effects exceeded SIIPS: SIIPS-NPS was negative for rating × HRF (t(7) = -2.713, p = 0.03008) and for event × HRF (t(7) = -3.432, p = 0.01096).

Since all days’ data also includes the training pain sessions, some perturbations in the association between pain and pain-decoder expressions could be expected. We therefore also ran a separate pre-training only analysis to validate the previous dataset. In the pre-training analyses effects were weaker and less consistent, as could be expected from a more restricted data pool.

NPS showed a marginally significant positive effect (t(7) = 2.366, p = 0.04986), whereas SIIPS did not show any effect (t(7) = 1.009, p = 0.3467). In direct paired comparisons on the pretraining day, SIIPS and NPS did not differ for either HRF model (rating × HRF: t(7) = -0.205, p = 0.843; event × HRF: t(7) = -1.051, p = 0.328).

Event-locked rating-decoder correlations confirmed the existence of temporal associations, peaking within the early post-onset window. For SIIPS, the correlation reached a significant peak at ≈2.8 s (lag 2 volumes) (t(7) = 5.033, q = 0.0106 BH FDR; Fig.3), whereas for NPS, the correlation showed a similar post-onset rise with the strongest effect at ≈4.2 s (lag 3 volumes) (t(7) = 3.850, q = 0.044; Fig. 3).

### 3.3 Up- and down-regulation of decoder during pain assessment

We further estimated run-level event-locked HRF-GLM beta coefficients separately for downregulation and upregulation trials and evaluated whether each condition differed from zero (one-sample t-tests) or whether upregulation differed from downregulation (Welch two-sample t-tests).

For the SIIPS decoder, none of the run-level estimates showed significant effects. Event-locked responses had negative mean values, but were not different from zero for both conditions (DOWN: t(9) = -0.73, p = 0.482; UP: t(9) = -0.06, p = 0.957, Fig. 4), and conditions weren’t different from each other (Welch t(17) = 0.426, p = 0.675).

**Fig. 4.**
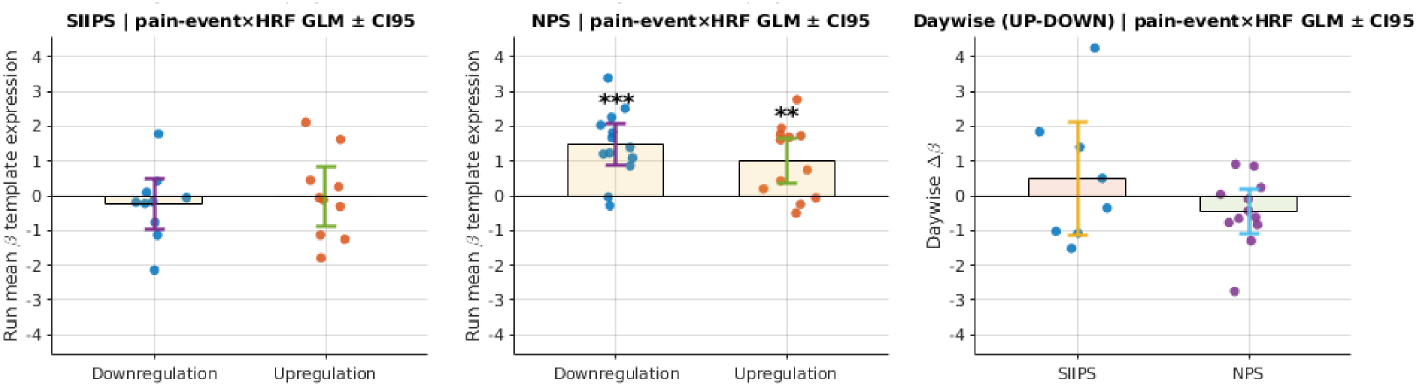
Pain encoding by the pain decoder activity (SIIPS or NPS) following different cue presentations. Columns 1 and 2 show how strongly pain responses predicted decoder expression separately for Upregulation and Downregulation trials. Column 3 shows the within-subject difference between conditions (Up - Down). We estimated these effects using pain-onset HRF-convolved GLMs applied to decoder time series that were z-scored relative to the distribution derived from resting-state data (see Methods). Models were fit separately for Upregulation and Downregulation trials within each run. To quantify regulation effects while accounting for repeated measurements, we first averaged run-level *β* estimates within each subject × day × condition. We then computed Δ*β* values (Column 3) by subtracting Downregulation from Upregulation estimates 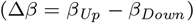 for each subject and day. Bars show the mean *β* estimates across runs for each condition. Dots indicate individual runs. Error bars denote 95% confidence intervals. Asterisks mark significant deviations from zero based on two-sided one-sample t-tests (*p < 0.05, **p < 0.01, ***p < 0.001)

For the NPS decoder, robust positive effects were observed as event-locked responses were strongly positive in both conditions (DOWN: t(12) = 5.33, p = 1.79 × 10-4; UP: t(11) = 3.35, p = 0.0065), but yet again without a significant UP–DOWN difference (t(23) = -1.14, p = 0.265).

Daywise contrasts further support run-wise data. For SIIPS, delta effect between Up- and Down-regulation runs was small and not different from zero for all three models (event-locked: t(7) = 0.73, p = 0.487; Fig. 4). For NPS, unexpectedly daywise contrasts were stably negative but non-significant (event-locked: t(11) = -1.58, p = 0.143). Thus, no statistically significant and consistent within-day modulation between up- and downregulation conditions was observed for any of the decoders.

### 3.4 Effects on pain ratings

We next examined whether decoded neurofeedback was accompanied by systematic changes in subjective pain ratings. All analyses were conducted at the day level, with a subject treated as a random factor. For each group (SIIPS vs. NPS cohorts), we first modeled day-mean ratings as a function of training (pre-vs. post-training) and condition (Up-vs. Down-regulation), then focused on within-day condition differences Δ(*condition*), and finally on baseline-corrected ΔΔ(*condition*) values that quantify how much the Up-Down difference changed relative to each subject’s own pre-training baseline.

For the SIIPS cohort, a mixed-effects model on day-level means (mean rating ∼ training phase × condition + (1|subject)) didn’t reveal any significant group or interaction effects (post-training effect: t(16) = -0.49, p = 0.63; upregulation effect: t(16) = 0.40, p = 0.70; interaction: t(16) = -0.79, p = 0.44; Fig. 5).

**Fig. 5.**
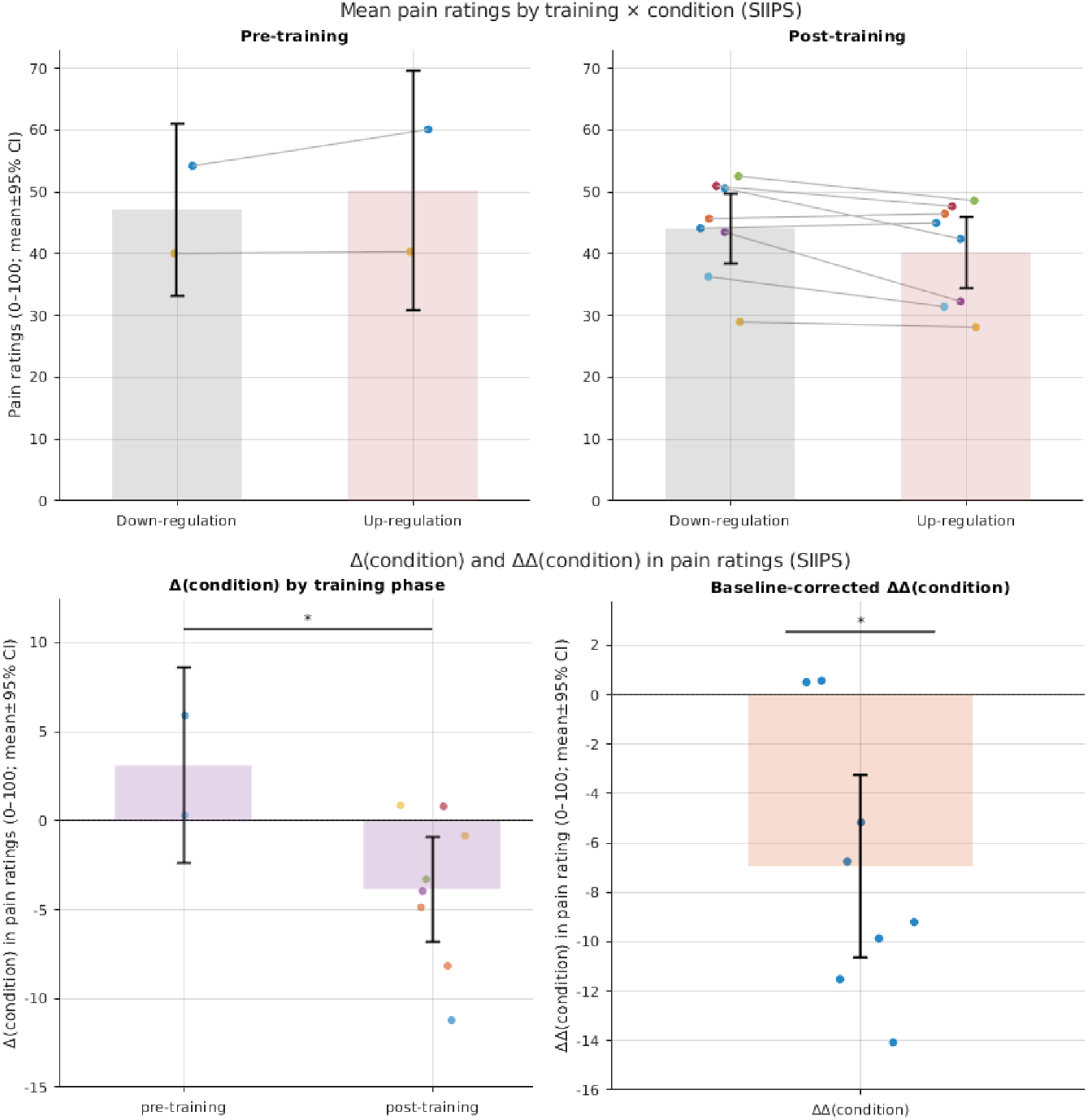
Effects of the SIIPS Up- and Down-regulation neurofeedback training on pain ratings. The top panel shows mean pain ratings (0–100 VAS units) for Upregulation and Downregulation trials before and after training. Values are averaged at the subject × day × condition level. Each colored dot represents one subject × day × condition mean. Grey lines connect the two conditions within the same subject and day. Each mean is based on 32 trials (8 ratings for each of 4 stimulus intensities; see Fig. 6 for intensity-wise results). The bottom left panel shows within-day condition differences, computed as Δ(*condition*) = rating(Up) - rating(Down), plotted separately for pre-training and post-training days. Each dot represents one subject × day value. We tested training effects using a mixed-effects model. Asterisks indicate a significant training effect (*p < 0.05). The bottom right panel shows baseline-corrected ΔΔ values for each post-training day. For each subject, we subtracted the pre-training Δ(*condition*) from the corresponding post-training Δ(*condition*): ΔΔ = Δ_*post*_ − Δ_*pre*_. Bars in the bottom panels indicate the mean Δ(*condition*) (or ΔΔ), and error bars show 95% confidence intervals across days (0–100 rating units).

However, both within-day (condition) model showed a significant training effect (post-vs pretraining t(8) = -2.32, p = 0.049), and baseline-corrected (condition) model showed a significant negative shift compared to 0 (t(7) = -3.67, p = 0.008; Fig. 5).

We therefore examined condition differences at each intensity level. For the within-day Δ(*condition*) measure (Up - Down) analysed separately by phase using, no level showed a significant effect after FDR correction in the pre-training data (all q≥0.26; Fig. 6). After training, however, Δ(*condition*) became strongly negative at intensity level3 (t(28)=-4.56, q=3.7×10^−4^), indicating substantially lower pain in the Up-cue trials than in Down-cue trials at this intermediate-high intensity. No other level showed reliable post-training effects (all q≥0.18).

**Fig. 6.**
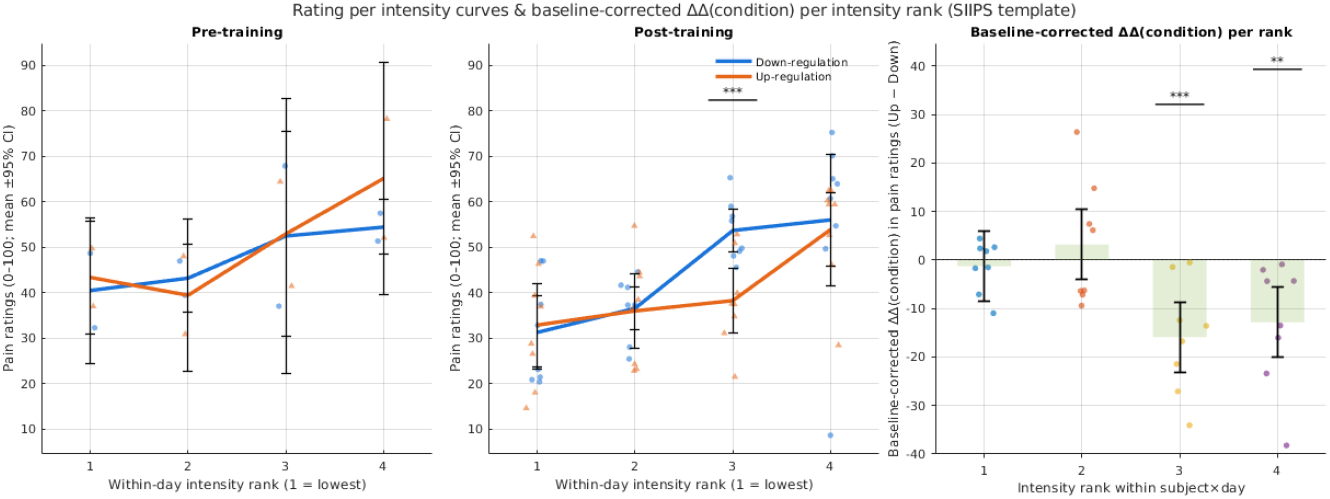
Intensity-wise pain effects of the SIIPS Up- and Down-regulation neurofeedback training on pain ratings. Left and middle panels: These panels show mean pain ratings (0–100 VAS) as a function of stimulus intensity rank (1 = lowest, 4 = highest temperature used for that subject). The left panel shows pre-training days, and the middle panel shows post-training days. Ratings are shown separately for Downregulation (blue) and Upregulation (orange) trials using the SIIPS decoder. Bars indicate the mean rating at each intensity rank. Error bars show 95% confidence intervals (0–100 rating units). Small symbols represent individual subject × day values. To test regulation effects at each intensity, we modeled within-day condition differences, defined as Δ(*condition*) = rating(Up) - rating(Down), using a linear mixed-effects model. Asterisks mark intensity ranks where the condition coefficient differed significantly from zero after Benjamini-Hochberg FDR correction across ranks (*q < 0.05, **q < 0.01, ***q < 0.001). Right panel: This panel shows baseline-corrected ΔΔ(*condition*) values at each intensity rank. For each subject and intensity, we computed ΔΔday = Δ_*post*_ − Δ_*pre*_, that is, the difference between the post-training and pre-training Δ(*condition*) values. Bars indicate the mean ΔΔ across subject × day units. Error bars show 95% confidence intervals. Dots represent individual subject × day values. Asterisks mark intensity ranks where the mixed-effects model coefficient differed significantly from zero after FDR correction across ranks (*q < 0.05, **q < 0.01, ***q < 0.001), indicating stronger post-training pain reduction for Upregulation trials at higher intensities.

Finally, baseline-corrected ΔΔ(*condition*) values (post-minus pre-training Δ(*condition*)) were analysed per level. Simple t-tests against zero showed a significant negative shift at level 3 (t(28)=-4.324, q(FDR)=7×10^−4^, Fig. 6) and rank 4 (t(28)=-3.490, q=0.003) with negligible effects at lower levels.

Thus, following SIIPS-based training, presentation of the cue associated with up-regulation of the SIIPS decoder expression was associated with relatively lower pain ratings than the down-regulation cue. Interestingly, these analyses suggest that SIIPS-linked cues preferentially reduced pain for higher-intensity stimuli, whereas downregulation cues did not produce the expected analgesic effect.

One possibility is that repeated engagement of SIIPS-related processes during upregulation training induced central habituation or reduced affective-motivational salience of high-intensity stimuli, leading to diminished subjective pain. Alternatively, upregulation attempts may have recruited higher-order evaluative or metacognitive control processes that subsequently attenuated pain perception, consistent with the proposed psychological nature of the SIIPS pattern.

These interpretations remain purely speculative and require confirmation in subject-level analyses. Furthermore, we must acknowledge that absolute means were noisy, and further confirmation on a subject-level analysis should be performed before making any conclusions.

For the NPS cohort, a significant main effect of phase was observed (post-training effect: t(33) = 3.61, p = 0.001), but not condition or interaction effects (upregulation effect: t(33) = -0.08, p = 0.94, interaction: t(33) = -0.52, p = 0.61; Fig. 7). Thus, this result suggests a general temporal effect rather than a condition-specific training effect.

**Fig. 7.**
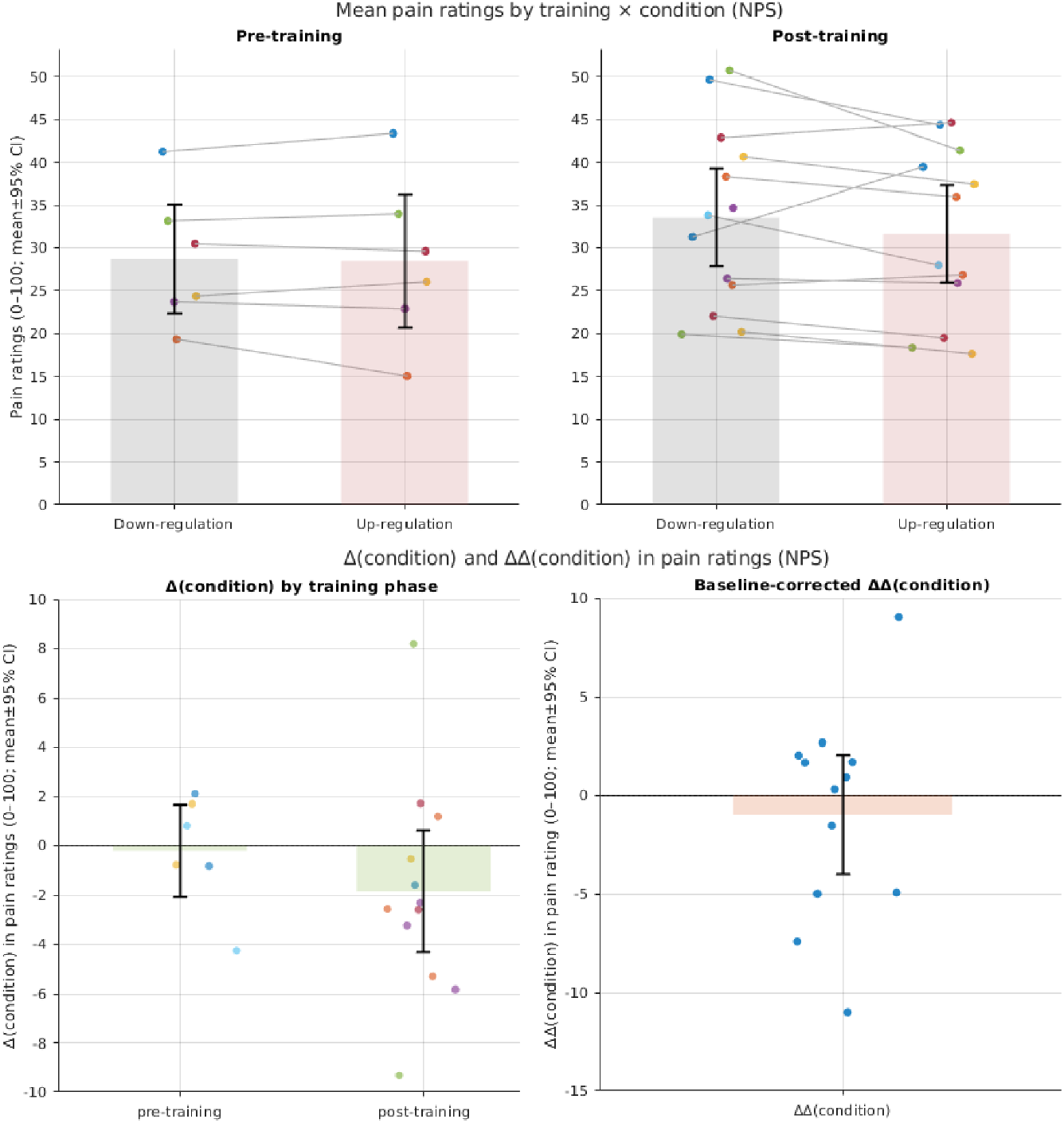
Effects of the NPS Up- and Down-regulation neurofeedback training on pain ratings. The top panel shows mean pain ratings (0–100 VAS units) for Upregulation and Downregulation trials before and after training. Values are averaged at the subject × day × condition level. Each colored dot represents one subject × day × condition mean. Grey lines connect the two conditions within the same subject and day. Each mean is based on 40 trials (10 ratings for each of 4 stimulus intensities; see Fig. 8 for intensity-wise results). The bottom left panel shows within-day condition differences, computed as Δ(*condition*) = rating(Up) - rating(Down), plotted separately for pre-training and post-training days. Each dot represents one subject × day value. We tested training effects using a mixed-effects model. Asterisks indicate a significant training effect (*p < 0.05). The bottom right panel shows baseline-corrected ΔΔ values for each post-training day. For each subject, we subtracted the pretraining Δ(*condition*) from the corresponding post-training Δ(*condition*): ΔΔ = Δ_*post*_ − Δ_*pre*_. Bars in the bottom panels indicate the mean Δ(*condition*) (or ΔΔ), and error bars show 95% confidence intervals across days (0–100 rating units).

Similarly, the within-day Δ(*condition*) model (post-vs. pretraining t(16) = - 0.90, p = 0.38) and baseline-corrected mode ΔΔ(*condition*) failed to achieve any significance (t(11) = -0.63, p = 0.54 vs. zero; Fig. 7).

For the NPS cohort, the same intensity mixed-effects model estimating phase, condition, intensity rank, and their interaction effects on mean ratings showed a strong main effect of intensity rank (F(1,140)=51.58, p=3.7×10^−11^), but no reliable main or interaction effects involving condition or training. Thus, compared to SIIPS data, NPS ratings are more valid and scale properly with intensity, but this dependence was similarly not detectably modulated by the Up vs. Down cues using this model and parameters.

Phase-specific linear models estimating intensity effects on Δ(*condition*) did reveal that, before training, Up-cue trials were slightly more painful than Down-cue trials at the highest intensity rank (t(20)=3.26, q=0.0157; Fig. 8), with no effects at lower ranks. But the effect has disappeared in the post-training, and no other pair-wise effects were observed.

**Fig. 8.**
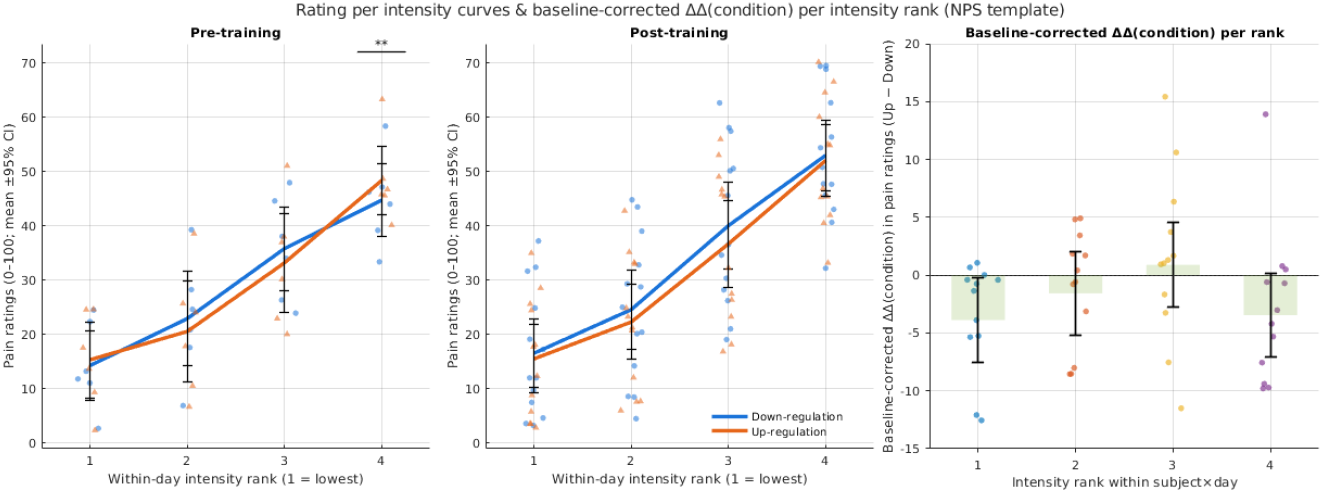
Intensity-wise pain effects of the NPS Up- and Down-regulation neurofeedback training on pain ratings. Left and middle panels: These panels show mean pain ratings (0–100 VAS) as a function of stimulus intensity rank (1 = lowest, 4 = highest temperature used for that subject). The left panel shows pre-training days, and the middle panel shows post-training days. Ratings are shown separately for Downregulation (blue) and Upregulation (orange) trials using the NPS decoder. Bars indicate the mean rating at each intensity rank. Error bars show 95% confidence intervals (0–100 rating units). Small symbols represent individual subject × day values. To test regulation effects at each intensity, we modeled within-day condition differences, defined as Δ(*condition*) = rating(Up) - rating(Down), using a linear mixed-effects model. Asterisks mark intensity ranks where the condition coefficient differed significantly from zero after Benjamini-Hochberg FDR correction across ranks (*q < 0.05, **q < 0.01, ***q < 0.001). Right panel: This panel shows baseline-corrected ΔΔ(*condition*) values at each intensity rank. For each subject and intensity, we computed ΔΔday = Δ_*post*_ − Δ_*pre*_, that is, the difference between the post-training and pre-training Δ(*condition*) values. Bars indicate the mean ΔΔ across subject × day units. Error bars show 95% confidence intervals. Dots represent individual subject × day values. Asterisks mark intensity ranks where the mixed-effects model coefficient differed significantly from zero after FDR correction across ranks (*q < 0.05, **q < 0.01, ***q < 0.001), indicating stronger post-training pain reduction for Upregulation trials at higher intensities.

Finally, baseline-corrected ΔΔ(*condition*), rank-wise t-tests also didn’t support the existence of any intensity-wise pain effects with rank 1 effect not surviving BH-FDR correction (rank 1: t(11)=-2.09, p=0.04, q=0.14; Fig. 8).

Overall, NPS training provides no evidence for a reliable Up-regulation vs. Down-regulation effect of perceived pain in this cohort. Similarly, the NPS cohort showed no consistent intensity-dependent effect of decoded neurofeedback on pain ratings.

## 4 Preregistration

With few exceptions [31], preregistered studies are still rare in the fMRI neurofeedback literature, limiting the field’s ability to distinguish reliable effects from context-specific artefacts [12, 32]. To address this gap, here we preregister a follow-up study, specifying key hypotheses and outlining experimental design, and analysis protocols.

The present study proposes to utilize a protocol closely aligned with the NPS group design, which, despite its limitations, seems to provide more consistent and expected results than the SIIPS design. Several considerations motivated this choice.

First, NPS group yielded the most reliable pain encoding by the Neurologic Pain Signature (NPS), demonstrating robust coupling between pain events and NPS expression across subjects. The existence of a clear association between the target cognitive pattern and the neurofeedback decoder is an important validation step that must be taken into account.

Second, the NPS group protocol employs a clear temporal separation between neurofeedback and pain direction (Up- or down-) regulation periods and trials. This separation reduces ambiguity about whether observed neural effects reflect long-term pattern induction, acute effects, pain processing, or their interaction.

Finally, we have chosen to keep the electrical stimulation that seems to have some clear ethical benefits compared to the thermal one, including shorter duration and less heat accumulation. In our setup, it also provides greater temporal control and an overall more stable setup.

Our neurofeedback trial design is kept the same as the one presented in the pilot study (Fig. 1) as it proved to be effective among the groups. Since we would like to additionally implement some other exploratory fMRI analysis of brain activity during the pain assessment trials, we are specifically opting here for the trials with varying event lengths to allow for better event separability. Our final trial design has the same structure as Fig. 1 and includes:

- Fixed time of a cue presentation before pain - 1.5s
- Additional jitter 0-2.8s
- Pain stimulation - 0.9s
- Jittered waiting time before the response 0-2.8s
- Response 4s
- Jittered inter-trial interval 0-2.8s

In addition to the inclusion criteria listed in the pilot study, we would also exclude subjects who failed to attend fMRI sessions at least once every 10 days. All our other methods planned to be used in the pre-registered study will also replicate the NPS group pilot design.

Here, we successfully reported subjects bidirectionally modulating their brain decoder expressions both for SIIPS and NPS, as well as some promising data on subjects’ potential ability to modulate their expression during pain stimuli presentation for NPS. However, translating these outcomes into behavioral changes proved to be challenging.

Our follow-up study will focus exactly on that, with the primary hypothesis being centered around behavioral outcomes, whereas the rest would be brain expression-related confirmatory analyses. Consequently, our stopping criterion is also defined based on the behavioral parameter. Given the pilot results, we also don’t propose any specific effect direction for the pain-related outcomes as now we expect that some complex interactions might exist (e.g,. pain decrease following decoder upregulation due to desensitization). None of the data presented here will be included in the final study for testing the hypotheses below.

### Primary Hypothesis (H1): Training alters the effect of cue presentation (Up-vs. Down-regulation) on pain ratings

Based on the pilot data, our primary chosen outcome is a subject-wise scalar effect ΔΔ(*condition*), defined as the subject mean baseline-corrected difference in condition effects between post-training and pre-training phases, computed as:

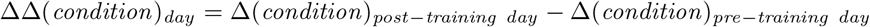

where Δ(*condition*) is calculated as the within-day difference between the two conditions for the subject:

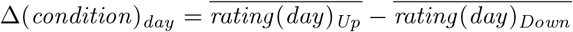

This parameter follows closely our statistics on the pain ratings reported here but additionally collapses the ΔΔ(*condition*) subject-wise, to appropriately account for the replicates biological nature.

Group-level evidence for deviation from zero will be evaluated using a one-sample t-test against zero (parametric) or an exact sign-flip test against zero (randomization-based; robust to non-normality).

Additionally, intensity-wise analyses will be performed in the same manner to assess any potential pain-intensity-dependent effects by calculating Δ(*condition*) and ΔΔ(*condition*) within the day for each intensity level separately.

Prior to primary hypothesis testing, distributional diagnostics will be performed on pain ratings (including inspection of residual variance, heteroskedasticity, and temporal autocorrelation). If substantial deviations from homoscedasticity or independence are observed, ratings will be transformed using either day-wise z-scoring or first-order autoregressive residualization. The decision rule will be based on data diagnostics, not on hypothesis outcomes. The selected normalization approach will then be applied uniformly across all subjects and conditions.

### Secondary Hypothesis (H2): Training alters effect of pain presentation on brain NPS decoder expression depending on condition (Up-vs. Down-regulation)

For each subject, pain-related BOLD responses will be modeled using a general linear model (GLM), preprocessed, voxel-wise z-score normalized, Neurologic Pain Signature (NPS) expression will be estimated as a dot product between voxel-wise activation estimates and the NPS weight map. Finally, the obtained score will be further normalized relative to the distribution of NPS scores on the resting state session of the same day. Pain onset regressors will be convolved with a canonical hemodynamic response function (HRF). Separate regressors will be defined for Up- and Down-regulation trials. Standard nuisance regressors (e.g., run intercepts) will be included. No orthogonalization between condition regressors will be applied.

The GLM will be fit to the NPS expression time series, yielding condition-specific regression coefficients (*β*_*Up*_ and *β*_*Down*_) that quantify the pain-locked modulation of NPS expression under each regulation condition.

For each day, a condition contrast will be computed as:

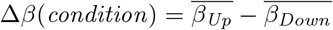

Then, the subject-wise training effect will be defined as the difference between post- and pre-training day contrasts:

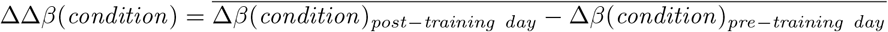

Group-level inference will be performed on subject-wise ΔΔ*β*(*condition*) values using a two-sided one-sample t-test against zero. Results will be reported as the mean ΔΔ*β*(*condition*) and its associated 95% confidence interval. No assumptions about effect directionality are made.

### Tertiary Hypothesis (H3): Training alters NPS decoder expression in neurofeedback trials depending on condition (Up-vs. Down-regulation)

Briefly, for each subject, pain-related BOLD responses will be modeled using a general linear model (GLM), preprocessed, voxel-wise z-score normalized, and Neurologic Pain Signature (NPS) expression will be estimated as a dot product between voxel-wise activation estimates and the NPS weight map. Finally, the obtained score will be further normalized relative to the distribution of NPS scores on the resting state session of the same day.

Neurofeedback-related NPS modulation will be further quantified using a predefined hemodynamically-shifted time window relative to the onset of each neurofeedback regulation period (i.e., applying a fixed lag to account for the HRF). Within each run, NPS scores will be averaged within this analysis window separately for Up-regulation and Down-regulation runs:

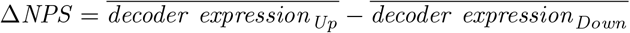

The mean subject Δ*NPS* value among all the runs will be used for further analyses. Group-level inference will be performed on subject-wise Δ*NPS* values using a twosided one-sample t-test against zero. Results will be reported as the mean Δ*NPS* and its 95% confidence interval. We expect Δ*NPS* to be > 0, thus reflecting an increase in NPS expression following cue presentation in Up-regulation runs compared to Down-regulation runs.

### 4.1 Sample size determination and stopping rule

For each run, condition-specific parameter estimates will be projected onto the predefined neural decoder, yielding decoder expression values for Up and Down conditions. A within-run contrast will be computed as Up - Down. These contrasts will be averaged across runs separately for pre-training and post-training sessions, yielding one pre- and one post-training value per subject.

The subject-wise change score (ΔΔ) will be defined as the post-training minus pretraining contrast. Group-level inference will be conducted on subject-wise ΔΔ values using a two-sided one-sample t-test against zero, and results will be reported as the mean ΔΔ with a 95% confidence interval. No effect direction is assumed.

All primary and secondary inferences treat subjects as the unit of analysis. Day-level and intensity-rank-level values are used for aggregation and descriptive visualization only and are not treated as independent observations.

Accordingly, the proposed study is designed around a precision-based stopping rule: data collection will proceed until the two-sided 95% CI half-width on the group mean ΔΔ(*condition*) is less than or equal to 2 rating units, or until a hard cap of 20 evaluable subjects is reached, whichever occurs first.

Across the available pilot subjects from the NPS cohort, the observed between-subject standard deviation of ΔΔ(*condition*) was approximately 4.5 rating units, yielding a 95% confidence interval (CI) half-width of approximately 3.2 units at n=6. Under the assumption that this variability remains approximately stable with additional data, the expected 95% CI half-width at sample size n scales as 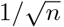, implying an expected half-width of approximately 2 rating units at n≈20. The value of 2 units was selected as a practically interpretable precision target that is feasible within the planned maximum sample size.

This stopping rule is non-directional and does not depend on statistical significance; instead, it ensures that the group-level effect is estimated with a predefined and interpretable level of precision. If the precision criterion (95% CI half-width ≤ 2) is met before N=20, the primary endpoint will be considered fully estimated. Recruitment may continue up to N=20 to support pre-specified secondary analyses (e.g., moderator and robustness analyses). The primary endpoint will be reported both at the time the precision criterion is met and at N=20 as a planned sensitivity update; interpretation will prioritize estimation (effect size and CI), not hypothesis testing.

If the CI half-width remains greater than 2 units at n=20, data collection will nevertheless stop, and the effect will be reported as imprecisely estimated within the available sample, consistent with the resource constraints and the observed heterogeneity in the pilot data.

All statistical tests, exclusion criteria, and stopping rules for confirmatory analyses are specified here prior to unblinding the main outcome data. Pilot results are used solely to assess feasibility and refine analytic sensitivity.

## 5 Acknowledgments

The study is supported by the Institute for Basic Science (IBS-R015-D2) in the Republic of Korea.

## 6 CRediT authorship statement

KD: Data curation, Formal Analysis, Investigation, Methodology, Project administration, Software, Visualization, Writing – original draft, Writing – review & editing; JSH: Investigation; WC: Investigation; DK: Resources; WW: Methodology, Resources, Software, Writing – review & editing; HL: Conceptualization, Funding acquisition, Methodology, Project administration, Resources, Supervision, Writing – review & editing; VT: Conceptualization, Methodology, Software, Writing – review & editing

## Notes

### Competing Interest Statement

The authors have declared no competing interest.

