## Supplementary Figure 1 for "Bidirectional modulation of pain by neurofeedback: Preliminary findings with fMRI at 7T"

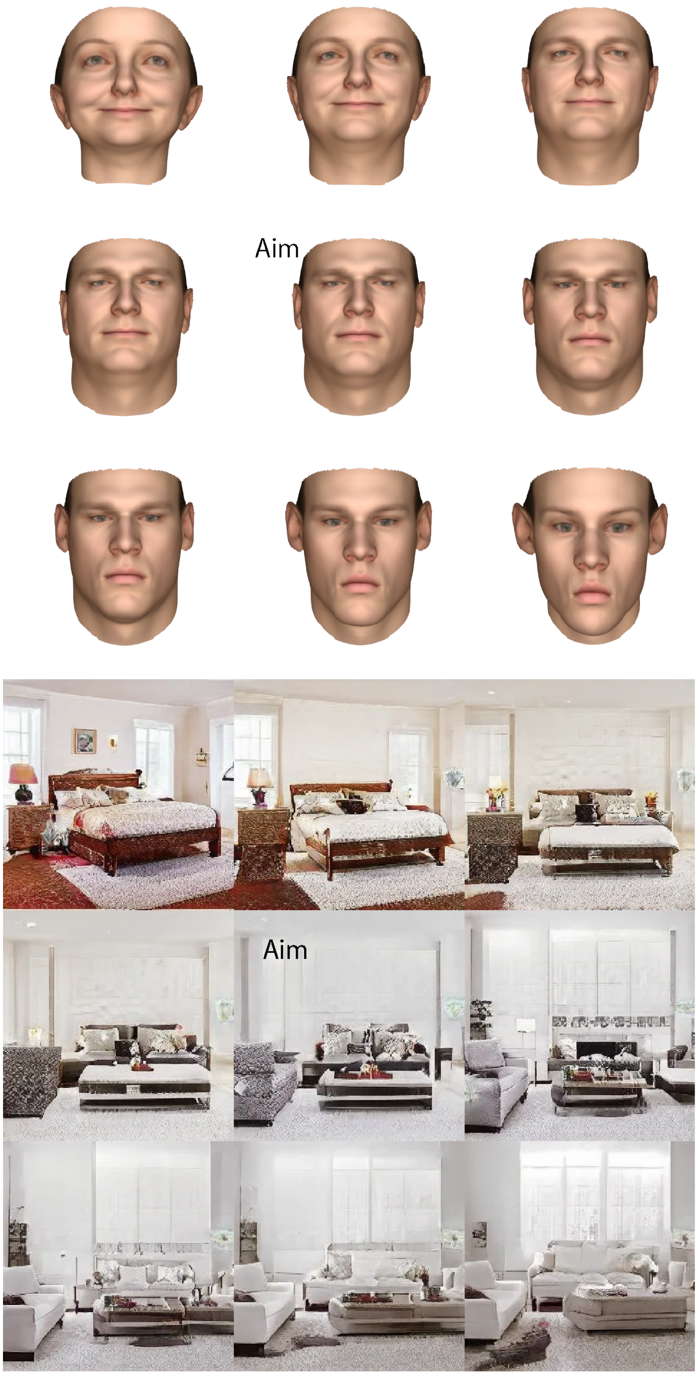


**Fig. S1** Main cues (central images labeled as aims) and distractors (the rest) used in the study.
